# Effects of unilateral cortical resection of the visual cortex on bilateral human white matter

**DOI:** 10.1101/689778

**Authors:** Anne Margarette S. Maallo, Erez Freud, Tina Tong Liu, Christina Patterson, Marlene Behrmann

## Abstract

Children with unilateral resections of ventral occipito-temporal cortex (VOTC) typically do not evince visual perceptual impairments, even when relatively large swathes of VOTC are resected. In search of possible explanations for this behavioral competence, we evaluated white matter microstructure and connectivity in eight pediatric epilepsy patients following unilateral cortical resection and 15 age-matched controls. To uncover both local and broader resection-induced effects, we analyzed tractography data using two complementary approaches. First, the microstructural properties were measured in the inferior longitudinal and the inferior fronto-occipital fasciculi, the major VOTC association tracts. Group differences were only evident in the ipsilesional, and not in the contralesional, hemisphere, and single-subject analyses revealed that these differences were limited to the site of the resection. Second, graph theory was used to characterize the connectivity of the contralesional occipito-temporal regions. There were no changes to the network properties in patients with left VOTC resections nor in patients with resections outside the VOTC, but altered network efficiency was observed in two cases with right VOTC resections. These results suggest that, in many, although perhaps not all, cases of unilateral VOTC resections in childhood, the white matter integrity in the preserved contralesional hemisphere along with residual neural activity might be sufficient for normal visual perception.

**Highlights:**

- There is well-circumscribed white matter damage in pediatric epilepsy after surgery
- White matter pathways are normal distal as well as contralesional to the resection
- Contralesional network properties differ after left or right hemisphere resection
- Preserved cortex and white matter may be sufficient for normal perception

## 1 Introduction

Recent studies have revealed that visual perception, for example contour integration and face and word recognition, is typically intact in children who have undergone unilateral visual cortical resection to manage pharmacoresistant epilepsy. One such case, UD, with surgical removal of the ventral occipito-temporal cortex (VOTC) of the right hemisphere (RH), showed normal intermediate and higher-order visual perceptual behavior despite the persistent hemianopia (Liu *et al*., 2018). This behavior was especially striking given that, as revealed through a longitudinal functional magnetic resonance imaging (fMRI) study, the resection impacted some category-selective regions in the RH (e.g. right fusiform face area) that are crucial for perceptual function. The findings of normal visual perception were corroborated in a study with a larger group of children, with the exception of two children with neurological comorbidities (Liu, Freud *et al*., 2019). Also, as was true for UD, too, normal category-selectivity in response to visual stimuli (e.g. images of faces and words) was observed in the remaining cortex (primarily in the VOTC of the structurally intact hemisphere) using fMRI.

A compelling question concerns the neural basis of the intact perceptual abilities in these patients. Given that surgery has been shown to be successful in managing the seizure disorder in the majority of the cases (Helmstaedter *et al*., 2019), it is important to understand the consequences and functional outcomes of this surgical procedure. One candidate mechanism that might impact behavioral outcome is the status of cortical connectivity postoperatively. Several studies have examined functional connectivity in pediatric epilepsy patients using graph-theoretic measures. For example, resting state fMRI studies have revealed that network modularity, path length, and global efficiency were independently associated with epilepsy duration (Paldino *et al*., 2017c). More relevant for establishing cognitive outcome after surgery, these same network measures have been associated with full scale IQ in children (Paldino *et al*., 2017b, 2019; Zhang *et al*., 2018). Of note is that network properties not only provide biologically and psychologically relevant measures in the pediatric epileptic brain cross-sectionally, but are also highly replicable longitudinally (Paldino *et al*., 2017a).

An alternative way to examine cortical connectivity is to focus on structure and quantify the integrity of the underlying white matter (WM) obtained from diffusion MRI. Some diffusion neuroimaging studies have revealed abnormalities even beyond the epileptogenic zone, including in regions that are typically associated with the default mode network (DeSalvo *et al*., 2014) and in the limbic system (Bonilha *et al*., 2012). In addition to insights that can be gleaned about the structural connectivity, diffusion MRI can also be used to assess the microstructural properties of the WM. For example, Pustina et al. (2014) revealed a decrease in fractional anisotropy (FA), which is a measure of WM integrity, in numerous tracts of patients with cortical resections, but especially in the ipsilesional tracts including the uncinate and fornix. However, these structural neuroimaging studies were conducted mostly with adults where the opportunity for plasticity is less than in younger patients.

Here, we used diffusion MRI with children who had undergone unilateral cortical resection. To our knowledge, there has not been a systematic assessment of WM in the VOTC postoperatively, and this is especially warranted in light of the previous findings of normal visual perception as described above. Moreover, because a reduction in structural connectivity is associated with reduced regional grey matter volume (Bonilha *et al*., 2010), a precise characterization of WM changes might be especially revelatory.

We adopted two different approaches to characterize the integrity of WM and its organization: first, we examined the microstructural properties of specific WM tracts that traverse the VOTC, and second, we evaluated cortical connectivity of the intact VOTC. In the first approach, we examined the local effects of resection on two VOTC WM tracts: the inferior longitudinal (ILF) and inferior fronto-occipital fasciculi (IFOF). We measured the FA, as well as the axial diffusivity (AD), a measure of diffusion parallel to the length of a tract, and the radial diffusivity (RD), a measure of the diffusion perpendicular to the tract. We selected these dependent measures as they generally offer clearer insight into the nature of injury than other measures, say, the mean diffusivity (Aung *et al*., 2013). We expected to observe hemispheric asymmetries in microstructural WM properties that could result from either an enhancement in the contralesional tracts, indicative of a compensatory mechanism, or an asymmetry in which compromised ipsilesional tracts are evident while the microstructural indices of contralesional tracts do not differ from those of the controls. In the second approach, to assess any possible WM changes on a broader scale, we used graph theory to characterize the network organization of the structurally intact contralesional VOTC and compared the network properties in the patients versus the control children.

The findings from this study have the potential to uncover important alterations in neural mechanisms following an assault on cortical development during childhood. Major changes to the WM might point to adaptive compensatory mechanisms, thus enabling normal perception. Conversely, the absence of significant alterations supports the hypothesis that the residual existing structure may be sufficient to support perception.

## 2 Materials and Methods

The procedures were reviewed and approved by the Institutional Review Boards of Carnegie Mellon University and the University of Pittsburgh. Parents provided informed consent and minor participants gave assent prior to the scanning session.

### 2.1 Participants

Participants included six pediatric patients with resections to the VOTC and two patients with resections outside the VOTC, and 15 age-matched typically-developing controls (three female, 12 male, mean age 14.5 ± 3.1 years; no age difference across groups, Wilcoxon rank sum test p>0.67). In total, three patients had resections to the RH and five had resections to the left hemisphere (LH). In seven of the eight patients, surgery was performed to manage pharmacoresistant epilepsy and, in the remaining case, surgery was performed for an emergent evacuation of cerebral hematoma at day one of life (Table 1).

**Table 1.**
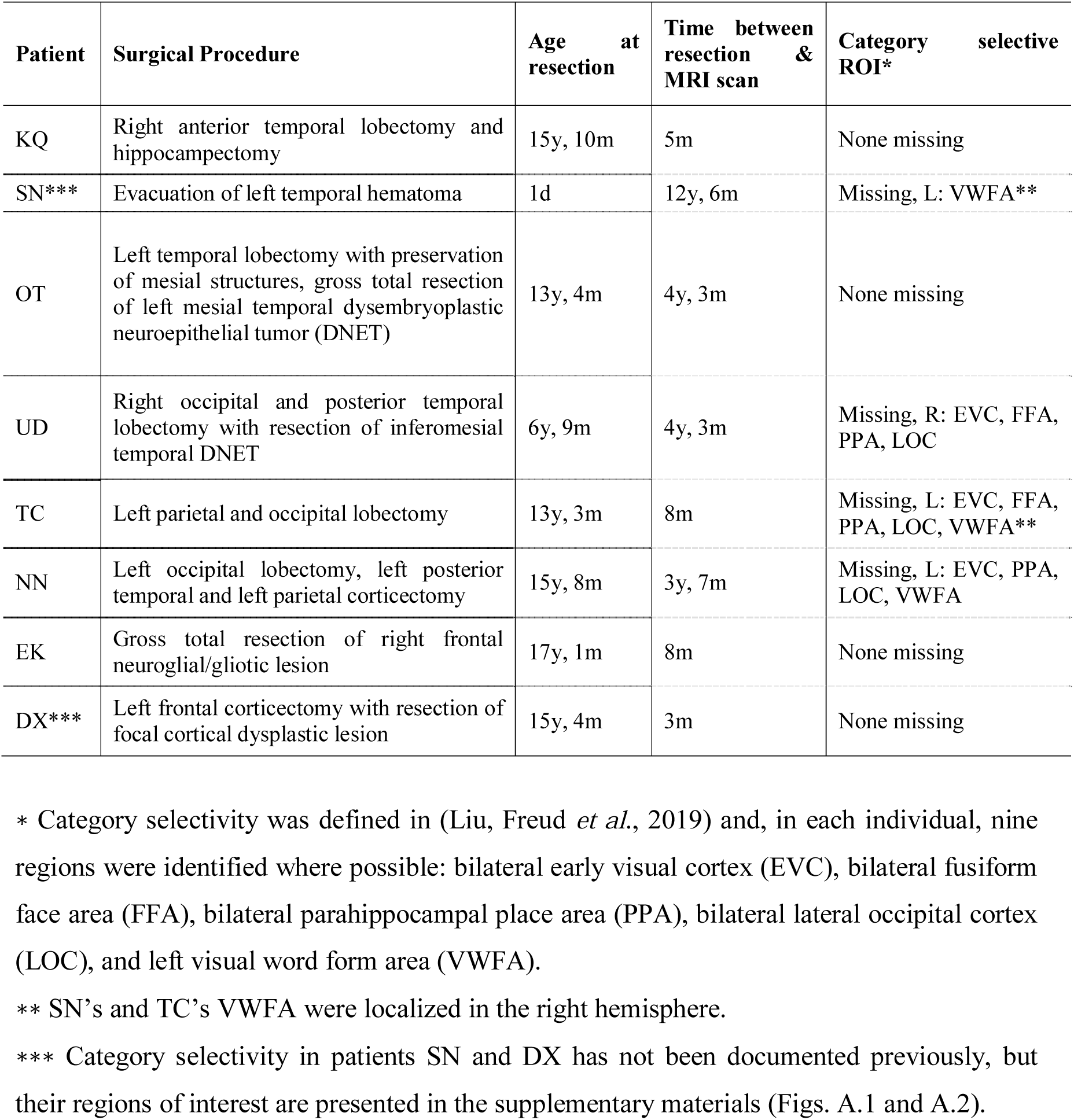
Surgery and data acquisition history of patients.

In six of these eight patients, the functional category-selective neural responses have previously been documented (Liu *et al*., 2018, Liu, Freud *et al*., 2019), but additional data from patients not included in those studies (SN and DX) are included in the Supplementary Materials (Figs. A.1 and A.2). Briefly, each participant completed three runs of an fMRI scan in which neural responses to images (faces, words, house and objects, as well as scrambled images) were measured, and used to define category-selective regions, which preferentially responded to one of the visual categories. In two patients with LH resection (SN and TC), the visual word form area, which is necessary for processing of words and typically localized in the LH, was instead localized in the RH.

The behavioral profiles for six patients have also been previously documented (Liu *et al*., 2018, Liu, Freud *et al*., 2019), and the data from the two additional patients (SN and DX) are presented here. Briefly, all participants completed several tasks designed to assess their visual perception (see Supplementary Materials Section C for more details). In these tasks, images were presented centrally, and patients were free to make saccades. Participants completed two tasks assessing their global perception: one that evaluates contour integration and one that measures their ability to detect “swirl” in glass patterns. Two further tasks examined their pattern recognition: one designed to assess object perception and one to measure face recognition.

We compared patients’ behaviors (Table 2), in terms of collinearity thresholds, accuracy, and response times, to age-matched controls. Although it is difficult to know exactly what the patients’ phenomenological experience was like and self-report is often imprecise (as is true for other cases of field defects such as hemianopia or quadrantanopia), the patients’ performance was nevertheless comparable to that of matched typically developing children on a wide range of dependent measures, indicating objective similarities in perception. An impairment was revealed both for the global perception and object recognition tasks in patient NN (who had polymicrogyria and low IQ as a complicating comorbidity), and for the global perception task in patient DX who had non-VOTC resection.

**Table 2.** Behavioral measures in patients with unilateral cortical resections Only NN and DX showed significant differences in performance relative to controls. Values in bold are significantly different to controls at p<0.05.

| Patient | Intermediate-level vision |  |  | High-level vision |  |  |  |
| --- | --- | --- | --- | --- | --- | --- | --- |
|  | Contour integration |  | Glass pattern | CFMT-C |  | Object matching |  |
|  | Threshold | Threshold | Threshold | Upright faces | Inverted faces | Accuracy | RT |
| | ($\pm 0$ collinearity) | ($\pm 20$ collinearity) | | | | | |
| KQ | 68.45 | 74.21 | 35 | 90.00% | 78.33% | 99.00% | 1018.14 |
| SN | 67.27 | 80 | 33.33 | 95% | 73.30% | 95.00% | 825.7 |
| OT | 54.01 | 65.73 | 31.67 | 62.50% | 55.56% | 99.00% | 929.73 |
| UD | 51.96 | 76.88 | 25.83 | 83.33% | 68.33% | 91.00% | 1366.96 |
| TC | 66.12 | 77.27 | 45.83 | 83.33% | 46.67% | 89.00% | 1047.66 |
| NN | **75.95** | 78.63 | 25.83 | 80.56% | 45.83% | 93.00% | **1726.1** |
| EK | 54.01 | 75.51 | 54.17 | 88.33% | 81.67% | 100.00% | 1192.12 |
| DX | **77.27** | 80 | 36.67 | 100.00% | 93.00% | 96.00% | 1208.9 |
| Control mean | 56.45±5.05 | 74.04±3.53 | 39.4±7.74 | 88.22±11.55% | 70.36±11.79% | 94.86±3.08% | 869.03±225.65 |

### 2.2 MRI Protocols

Images were acquired on a Siemens Verio 3T scanner with a 32-channel head coil at Carnegie Mellon University. A T1-weighted image with 1×1×1mm3 resolution (MPRAGE, TE=1.97ms, TR=2300ms, acquisition time=5m21s) and one set of diffusion-weighted images with 2×2×2mm3 resolution (2D EPI, 113 directions, maximum b=4000s/mm2, TE=121ms, TR=3981ms, acquisition time=7m46s) were acquired for all participants. The neuroimaging data presented are being uploaded to KiltHub, Carnegie Mellon’s online data repository (reserved DOI: 10.1184/R1/9856205). However, the DOI will appear empty as the process is pending institutional approval. We trust that this will be available online after the review process.

### 2.3 Microstructural properties of VOTC white matter pathways

This first approach allowed us to determine whether there were local microstructural alterations to either of the two major WM tracts that traverse the VOTC (Catani *et al*., 2003, Catani, 2006). The ILF connects the extrastriate occipital lobe and the anterior temporal lobe, running along the lateral and inferior wall of the lateral ventricles. The IFOF is an indirect pathway that runs from the occipital to the temporal lobes and then passes through the uncinate and corona radiata along the lateral border of the caudate nucleus, finally terminating in the frontal lobes.

The diffusion data were reconstructed in DSI Studio (http://dsi-studio.labsolver.org) using generalized q-sampling imaging (Yeh *et al*., 2010) with 1.25 diffusion sampling ratio. We used a deterministic fiber tracking algorithm (Yeh *et al*., 2013) with whole-brain seeding, 0.20 quantitative anisotropy threshold, 0.20mm step size, 20mm minimum length, and 150mm maximum length to generate four sets of tractograms with one million streamlines each, at angular thresholds of 30°, 40°, 50°, and 60° that were ultimately combined to create one effective whole-brain tractogram. These thresholds were consistent with values that allow for robust fiber tract reconstructions as used in the literature (Dennis *et al*., 2015).

Next, we delineated the ILF and IFOF in the patient group, in both the ipsilesional and contralesional hemispheres, as well as in both hemispheres of the control group. We filtered the global tractogram described above to extract the tracts using the following standard procedure, which has been previously used successfully to generate reproducible results (Wakana *et al*., 2007). The ILF was defined using two regions of interest (ROIs) as inclusion masks. One ROI included the entire occipital cortex and was located coronally at the most anterior slice of the parieto-occipital sulcus. The other ROI included the entire temporal lobe and was located coronally at the most posterior slice in which the temporal lobe was detached from the frontal lobe. The IFOF was defined similarly: one ROI included the entire occipital cortex and was located coronally halfway between the anterior and posterior slices of the parieto-occipital sulcus. The other ROI included the entire frontal lobe and was located coronally at the anterior edge of the genu of the corpus callosum. For both tracts, streamlines that projected across the midline were discarded.

We measured the FA, AD, and RD of the tracts also in DSI Studio. These indices were only measured along the tracts, which were external to the resection areas (i.e. the tracts were located superior or inferior to, or more medially/laterally with respect to the resection). A lesion mask was manually drawn based on the anatomical image to enable us to determine the extent of the resection in the anteroposterior direction and, combined with anteroposterior profiles of the microstructural indices, we were able to determine the locations of any abnormalities relative to the resection.

We computed the mean microstructural indices in two regimes. In the first regime, we computed the mean indices over the entire length of each tract and used these values for between-group comparisons in patients and controls. In the second regime, we computed the mean indices over segments of each tract and used these values in single-subject analyses. Given the heterogeneity in the resection site in the patients and the generally large extent of the resections, we deemed it sufficient to bisect the tracts into two segments representing the anterior or the posterior segment of each tract. For instance, if a tract was of length *L*, we bisected it into segments, each of length *L*/2 and this was done independently for the left and right ILF and IFOF in each individual.

### 2.4 Network properties in the contralesional hemisphere

This second approach allowed us to determine whether there were broader resection-induced changes in the connectivity of regions in the occipito-temporal cortex. However, because of the heterogeneity in the location and extent of the resection and the variability in the preserved ipsilesional tissue, it was difficult to study the ipsilesional hemisphere across the patient group. Additionally, automated parcellation of the ipsilesional hemisphere was likely to be unreliable given the structural abnormalities and manual demarcation of regions may not have been sufficiently systematic across cases. Therefore, we studied only the contralesional hemisphere and its network properties.

For each participant, we used only a single tractogram to define the network connectivity, with an angular threshold of 60°. All other thresholds were the same as in Section 2.3, except for the number of streamlines. Instead of terminating tractography once a specific number of streamlines was reached, here, we controlled the seed density (20/voxel) by placing seeds only in the mask encompassing the preserved hemisphere in each individual to control for the variability in the participants’ head sizes.

Next, we used cortical ROIs based on the parcellation of the anatomical images using the Destrieux atlas (Destrieux *et al*., 2000) in FreeSurfer (Dale *et al*., 1999). The parcellation output was visually inspected and confirmed to be reliable in the structurally preserved hemisphere of the patients (see Supplementary Materials Section B for list of the regions comprising the VOTC that were used as network nodes). We generated a connectivity matrix in DSI Studio, such that, for any ROI pair, the connectivity value was the mean FA of all tracts that pass through both ROIs. We binarized the connectivity matrix for each individual by using a threshold of one standard deviation below the mean FA (excluding FA values of 0 – i.e. ROI pairs that were not connected by any streamline was not used in the computation of the threshold). Any value higher than the threshold was assigned a value of 1, and every other value was assigned a value of 0. With this binary connectivity matrix, we computed four standard network measures: transitivity, modularity, characteristic path length, and global network efficiency using the Brain Connectivity Toolbox (Rubinov and Sporns, 2010) for MatLab. We then compared these graph-theoretic measures between patients and controls using between-group permutation and single-subject analyses. Additionally, we also explored, on an individual level, the effects of using a different threshold in binarizing the connectivity matrices.

### 2.5 Statistical analyses

#### 2.5.1 Group comparisons

In the between-group comparisons, given the small number of VOTC resection patients, we generated 100,000 permutations of data from the two groups (six VOTC resection patients and 15 controls) and used a two-sample t-test, thus creating a null distribution of t-scores. We did not include the non-VOTC resection patients for this analysis. Then, p-values were computed as the ratio of values that were more extreme than the actual patient versus control group t-score relative to the number of permutations (the number of |t_permutation_|>t_actual_ divided by 100,000).

#### 2.5.2 Single-subject level comparisons

In the single subject analyses, we used established statistical procedures (Crawford and Howell, 1998) to compare each individual patient’s data to the data from the controls. With a normative sample size (degree of freedom=14, t_c_=2.145), a modified t-test was used in which each patient was treated as a sample of n=1, thereby eliminating the contribution of the single subject to the estimate of within-group variance.

## 3 Results

### 3.1 ILF and IFOF in VOTC resection patients

We defined the ILF and IFOF in six children with unilateral resections to their VOTC, as well as in the controls. We were able to reconstruct the ILF and the IFOF in most patients, even in the ipsilesional hemisphere, as long as the streamlines were outside the resection (Fig. 1), with the exception of the ipsilesional ILF and IFOF in KQ and the ipsilesional IFOF in SN.

**Figure 1:**
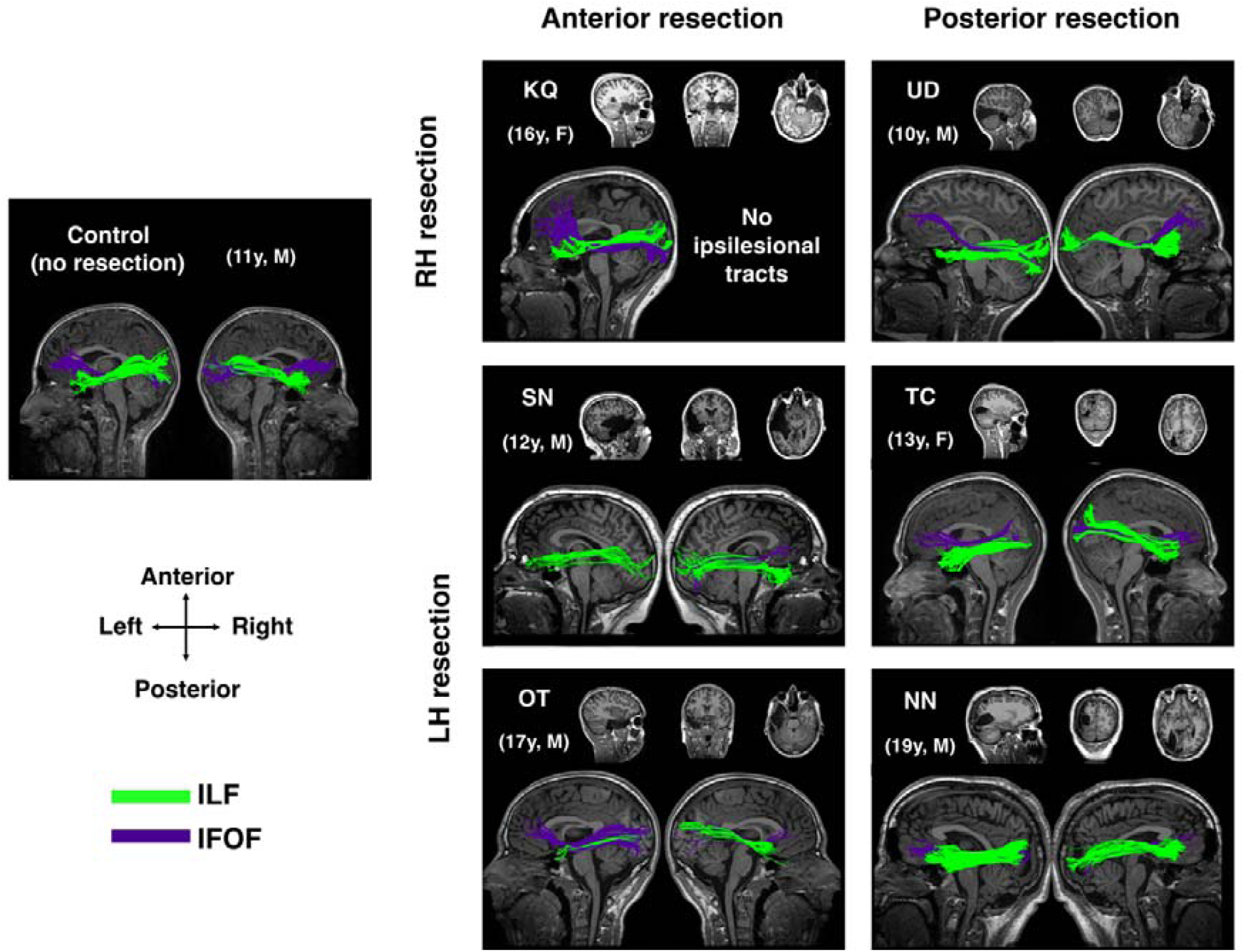
ILF and IFOF in patients with VOTC resections. ILF and IFOF in exemplar control and patients with VOTC resections to their right hemisphere (RH) and left hemisphere (LH). Anatomical images show the resection site in each patient. Both contralesional and ipsilesional tracts were present in most patients, except for right (ipsilesional) ILF and IFOF in KQ, and left (ipsilesional) IFOF in SN. Ages shown for patients are at time of data acquisition.

#### 3.1.1 Between-group differences in microstructural properties of the tracts

To evaluate the differences between the patients and controls, we computed the mean FA, AD, and RD in both of the identified tracts (Fig. 2). Perhaps surprisingly, there were multiple peaks in the null distribution histograms, a result of having extreme values from patients in the pooled data set from both groups that was resampled with no replacement. Relative to this derived null distribution, we found that, for the ipsilesional ILF, FA was lower and both AD and RD were higher in patients than controls (actual between group |t|>2.5, all p<0.01). Meanwhile, for the ipsilesional IFOF, there were only significant differences between patients and controls on RD (higher in patients than controls, between group t=2.5, p<0.05). In contrast, for both the contralesional ILF and IFOF, the findings did not differ significantly between the patients and the controls on any of the microstructural indices (between group |t|<1.69), all p>0.11). Moreover, the results were the same irrespective of whether the patient data were compared to data from either the LH or the RH of the controls, or to the mean of the bilateral tracts in controls.

**Figure 2:**
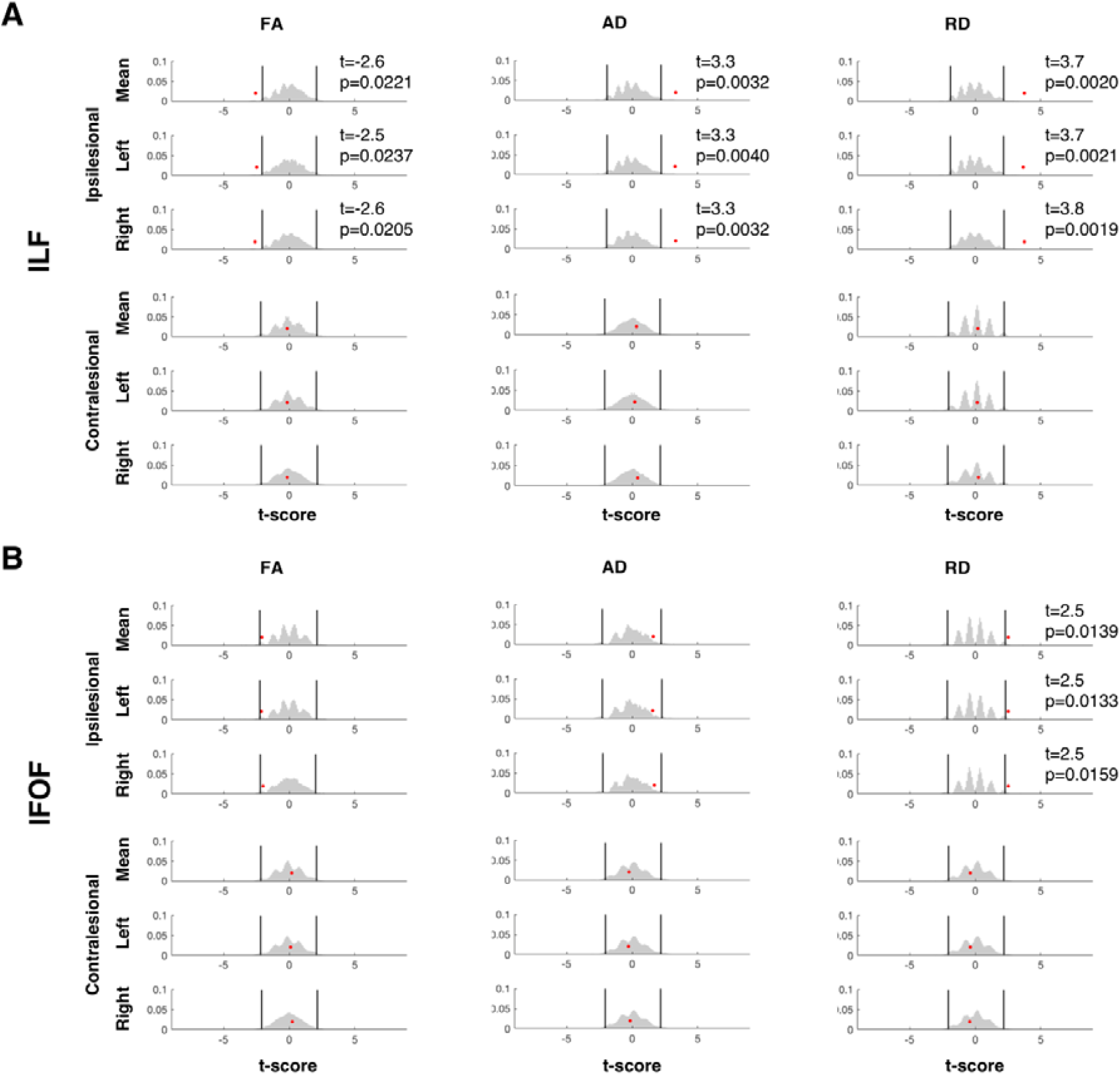
Group comparison of microstructural indices between VOTC resection patients and controls. Results of permutation testing of mean microstructural indices of the (A) ILF and (B) IFOF. There were significant group differences at α = 0.05 between controls and patients (red dots) for all measures in ipsilesional ILF but only in RD for the IFOF. Both contralesional tracts had group differences of p > 0.11 across all microstructural indices. Patient data were compared to the mean of the indices of both hemispheres in controls (top rows) or to indices from either hemisphere (2nd and 3rd rows). Black vertical lines indicate upper and lower 2.5th percentile of the null distribution (gray histogram).

#### 3.2.2 Microstructural properties of the ILF

Next, we wished to determine whether the observed ipsilesional group differences were present along the entire length of the tracts or restricted to the proximate site of the resection. We plotted the three different microstructural indices along the anteroposterior axis of the ILF (Fig. 3A). In an exemplar control participant (top rows, Fig. 3A), the respective microstructural indices of the ILF were each comparable bilaterally along the entire length of the tract. In patients in whom we recovered bilateral ILF, unlike in controls, there appeared to be differences in the microstructural indices of the two hemispheres: visual inspection suggests that FA was lower, and AD and RD were higher in the ipsilesional than in the contralesional ILF (consistent with the finding from the between-group analysis above). Furthermore, these differences appeared to be confined to the extent of the resection of each patient (preserved hemisphere serves as within-subject comparison, Fig. 3A). Thus, in patients SN and OT who have anterior temporal lobe resections, the within-subject hemispheric asymmetry was evident mostly in the anterior segment of the ILF. While in patients UD, TC, and NN, who had more posterior resections, the within-subject hemispheric asymmetries were evident in the posterior segment of the ILF.

**Figure 3:**
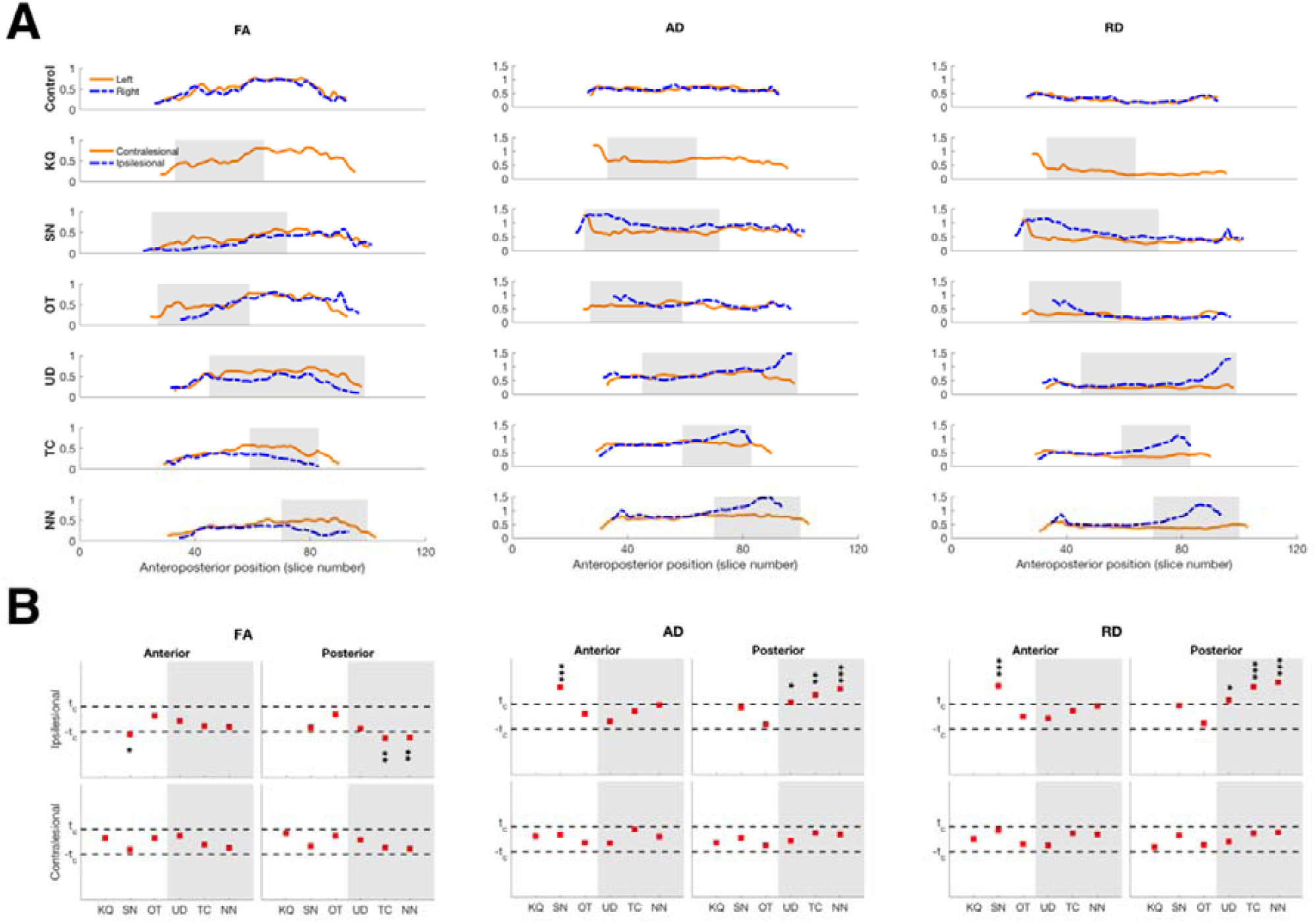
Microstructural indices of the ILF. (A). Anteroposterior profiles of FA, AD, and RD in an exemplar control and in patients with resections to their VOTC. Shaded regions indicate extent of resection. (B). Single-subject comparisons against normative group values (n=15) showed significant differences between ipsilesional and corresponding ILF in controls only for values in the segment corresponding to the resection in each patient. Shaded areas indicate patients with posterior resection, white areas indicate patients with anterior resection. p-values uncorrected for multiple comparisons: *p < 0.05,** p < 0.01,*** p < 0.001.

We quantified the microstructural damage by comparing the mean indices in both the anterior and posterior segments of the ILF in patients to the corresponding values from controls (Fig. 3B), and found that only patient OT exhibited normal microstructural indices across the board. Critically, for the other patients, the significant differences to the microstructural indices of the ILF was confined to the extent of the resection (Fig. 3B, all |t|>2.4, all p<0.05). That is, compared to controls, patients with posterior resections had compromised microstructural indices only in the posterior segment of the ILF and patient SN (the only patient who had an anterior resection with bilaterally defined ILF) exhibited compromised microstructural indices only in anterior microstructural indices of the ILF. Moreover, both segments of the contralesional ILF exhibited normal microstructural indices compared to controls.

#### 3.1.3 Microstructural properties of the IFOF

As with the ILF, the anteroposterior profiles of microstructural indices of the IFOF in patients are also different from those of controls, as well as from the profiles of the same measures in the ILF in patients. In an exemplar control participant (top rows, Fig. 4A), the respective microstructural indices of the IFOF were each comparable bilaterally along the entire length of the tract, similar to the ILF. And in patients in whom we recovered bilateral IFOF, visual inspection indicates that within-subject hemispheric differences in the IFOF were present only in patients with more posterior VOTC resections (preserved hemisphere serves as within-subject comparison, Fig. 4A), wherein the following patterns, similar to those from the ILF, were seen: FA was lower, AD and RD were higher in the ipsilesional than in the contralesional IFOF. However, the IFOF could not be defined in two of the three patients with anterior resection and so this finding ought to be interpreted with caution. Thus, in patients with posterior resections (UD, TC, and NN), the within-subject hemispheric asymmetries were evident in the posterior segment of the IFOF.

**Figure 4:**
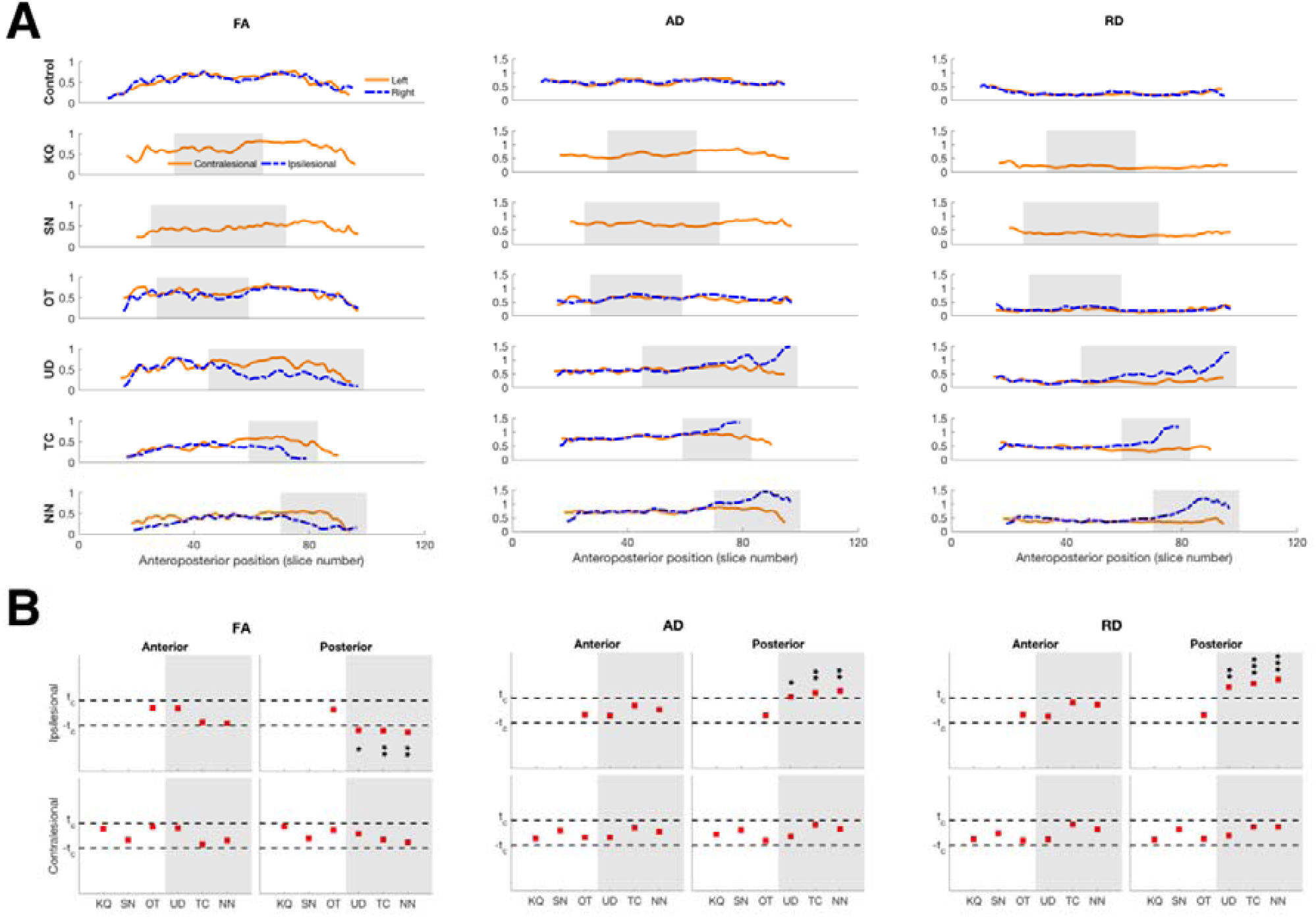
Microstructural indices of the IFOF. (A). Anteroposterior profiles of FA, AD, and RD in an exemplar control and in patients with resections to their VOTC. Shaded regions indicate extent of resection. (B). Single-subject comparisons against normative group values showed significant differences between ipsilesional and corresponding IFOF in controls only for values in patients with posterior resection. Shaded areas indicate patients with posterior resection, white areas indicate patients with anterior resection. p-values uncorrected for multiple comparisons: *p < 0.05,** p < 0.01,*** p < 0.001.

We also quantified the microstructural damage by comparing the mean indices in both the anterior and posterior segments of the IFOF in patients to the corresponding values from controls (Fig. 4B), and found that the significant differences to the microstructural indices of the IFOF was confined to the extent of the resection (Fig. 4B, all |t|>2.4, all p<0.05). That is, the patients with posterior resections had compromised microstructural indices only in the posterior segment of the IFOF. And similar to the ILF, the contralesional IFOF exhibited normal microstructural indices compared to controls.

#### 3.1.4 Normal microstructural properties of the ILF and IFOF in non-VOTC resection

We next characterized the ILF and IFOF (Fig. 5A) in the two patients with pharmacoresistant epilepsy who had resections outside their VOTC (EK and DX). Using identical procedures as in the children with VOTC resections, we found no qualitative within-subject hemispheric asymmetry in the anteroposterior profiles of the microstructural indices of either tract in these two patients (Fig. 5B-C) and visual inspection suggests that these plots resemble the profiles seen in controls (Figs. 3A and 4A, top rows). Furthermore, single-subject analysis revealed no significant differences in either case compared to the controls across any of the indices (all |t|<1.23, all p>0.24), be it in the ipsilesional or in the contralesional tracts.

**Figure 5:**
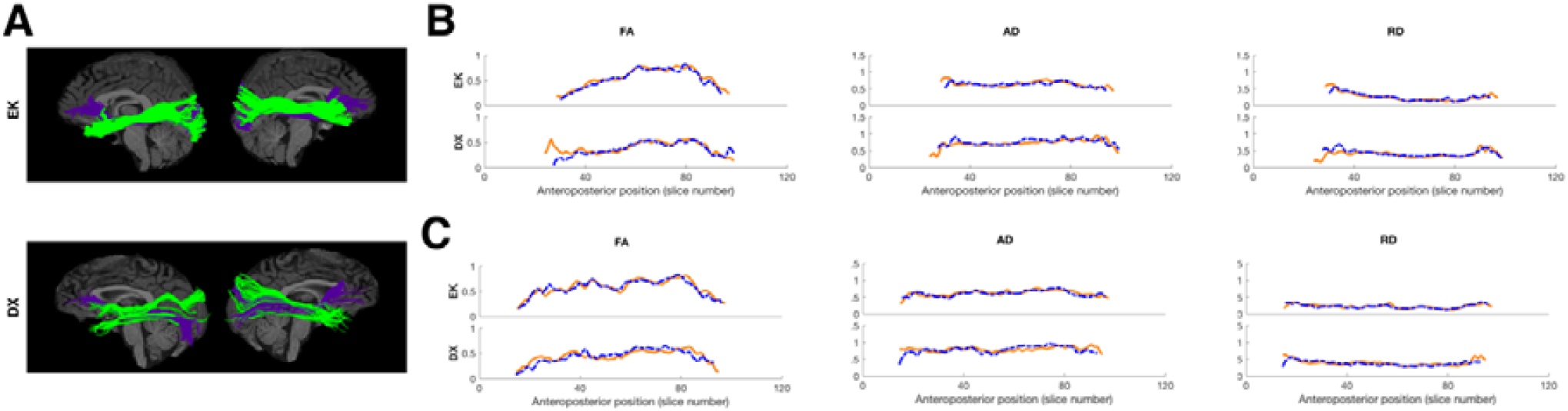
ILF and IFOF in children with resections outside VOTC. Microstructural indices of the (B) ILF and (C) IFOF in patients with resections outside their VOTC. There were no single-subject level differences to control values for any of the indices of bisected tracts, all |t|<1.23, all p>0.24. Same colors and orientation as in Fig. 1.

### 3.2 Network properties of the contralesional occipito-temporal cortex

Thus far, we have uncovered local changes to specific WM tracts by revealing microstructural damage restricted to the extent of resection. In a complementary approach, we wished to examine whether there were also broader resection-induced changes to the structural connectivity of the visual cortex. We used the same between-group and single-subject approaches to characterize the network properties in the intact contralesional hemisphere in the VOTC resection patients as in previous sections.

#### 3.2.1 Between-group comparisons of network properties

First, we combined the data from patients and controls in order to perform permutation testing on the network properties including transitivity, modularity, path length, and efficiency. On a group level (Fig. 6A), relative to the null distribution, there were no difference between patients and controls, and this was true for all network properties (all |t|<0.4568, all p>0.65). Note that now, there is only a single peak, in the histograms, as the patient group values fall within the same normal distribution as the controls.

**Figure 6:**
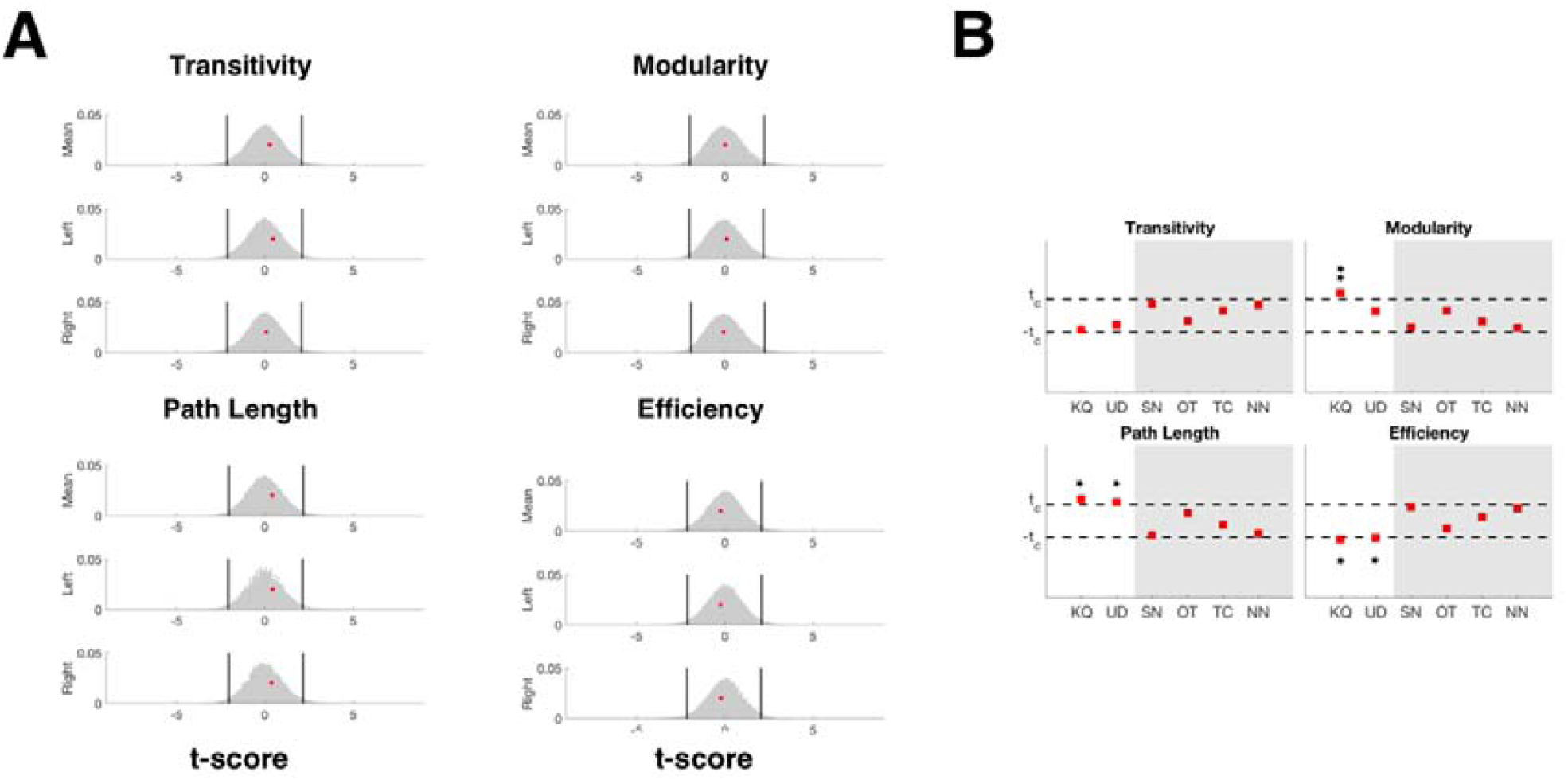
Comparison of graph-theoretic measures between VOTC resection patients and controls. (A) Results of permutation testing of network properties reveal normal measures in patients compared to controls on a group level (all p>0.65) and are consistent whether patient data were compared to the mean of both hemispheres in controls (top rows) or to either hemisphere (2nd and 3rd rows). (B) Patients with LH VOTC resections exhibit normal network properties while patients with RH VOTC resections (KQ and UD) exhibit significantly different network properties compared to controls. Shaded areas indicate patients with posterior resection, white areas indicate patients with anterior resection. p-values uncorrected for multiple comparisons: *p < 0.05,** p < 0.01.

#### 3.2.2 Single-subject comparison of network properties

Next, we looked at the network properties on a single-subject level compared to the normative group and we observed a difference in the network properties of a subset of the patients with VOTC resections compared with those of the controls. The four patients with LH VOTC resections had normal graph-theoretic values on all four dependent measures. The two patients with RH VOTC resections (KQ and UD), however, exhibited altered network properties, relative to the controls (Fig. 6B). KQ had higher modularity and characteristic path length and lower network efficiency (all |t|>2.4, all p<0.05), and UD had higher characteristic path length and lower network efficiency (both |t|>2.2, both p<0.05). Importantly, consistent with the results in Section 3.1.4, there were no changes to the network properties in the two patients with resections outside their VOTC (all |t|<1.94, all p>0.11).

#### 3.2.3 Effects of using different thresholds on the network properties

Network analysis is contingent on using robust connectivity matrices. Thus, we also explored the effects of using different thresholds in binarizing the connectivity matrices (see supplementary materials, Fig. D.1). At a higher threshold, there were no significant differences between any of the patients and controls in any of the dependent measures. At lower thresholds, only one patient with left resection, OT, had a significantly different path length and efficiency (all |t|>2.14, all p<0.05), and in the two patients with right resection, KQ had significantly different properties to controls, while these effects in UD disappeared.

## 4 Discussion

Surprisingly, in contrast with the visual perceptual impairments that ensue even after a fairly circumscribed lesion to VOTC in adulthood, visual perception in pediatric epilepsy patients with relatively large resections appears remarkably normal (as shown in Liu *et al*., 2018, Liu, Freud *et al*., 2019). The key question addressed here is whether alterations in WM underlie the positive visuoperceptual outcomes. As has already been documented (Taylor *et al*., 2015), behavioral impairments are not always correlated with the size and site of affected cortex, and aberrant cortico-cortical interactions might positively or negatively affect the expected brain-behavior correspondence. To elucidate possible neural mechanisms supporting the normal perceptual competence in the patients with unilateral resections presented here, we characterized the integrity of the two major WM association tracts that traverse the VOTC, the ILF and the IFOF, as well as the WM network properties.

### 4.1 Ipsilesional tracts were present in some patients

We first reconstructed the ILF and IFOF, which have previously been shown to underlie object recognition (Thomas *et al*., 2009; Tavor *et al*., 2014; Behrmann and Plaut, 2013; Decramer *et al*., 2019; Ortibus *et al*., 2011) and mediate visual perception (Mishkin *et al*., 1983). We were able to delineate all contralesional tracts and even the ipsilesional tracts in most patients, except in those with extremely large resections of the entire anterior temporal lobe. That we could not demarcate all ipsilesional tracts was not surprising given the extent of resection that affected the anterior temporal lobe in some of these patients. Moreover, the large resections caused automated tractography algorithms to fail in producing streamlines that survived the tractography thresholds, as voxels containing cerebrospinal fluid, which now filled the resection site, have different diffusion values to voxels containing WM. Nevertheless, the mere presence of the ipsilesional ILF and IFOF in most of the patients suggests that any damage caused by the removal of cortical tissue does not result in an obvious cascade of axonal injury that affects the entire ipsilesional hemisphere beyond the resection site.

#### 4.1.1 Microstructural changes were specific to site of resection

To provide a more quantitative analysis of the WM, we characterized the tracts in terms of their FA, AD, and RD. These indices were chosen because they offer a mechanistic relationship to the nature of injury (Aung *et al*., 2013); while FA has typically been used to characterize microstructural integrity, AD and RD have been shown to be more sensitive biomarkers for WM defects (Arfanakis *et al*., 2002; Xu *et al*., 2007; Moen *et al*., 2016), though the associated mechanisms might differ. Our results showed that, in children with a VOTC resection, there was damage to the ipsilesional ILF and IFOF that manifested in one or all of the following neuroimaging markers: there was within-subject hemispheric asymmetry in the microstructural indices – FA was lower, while both AD and RD were higher in the ipsilesional than in the contralesional tract. Importantly, the microstructural deficits in the pediatric epilepsy resection cases were only observed in regions proximal to the resection. In the ILF, abnormalities were evident in patients with either anterior or posterior resection, while in the IFOF, abnormalities were seen only in patients with posterior resections. This is unsurprising as the trajectory of the IFOF was more medial (Makris *et al*., 2007), while the anterior resections in the patients here were more lateral, thereby sparing the IFOF. Distal to the resection (either antero- or retrograde) in the ipsilesional tracts, the microstructural indices were normal compared to controls. Moreover, the microstructural indices in the contralesional ILF and IFOF in VOTC resection patients were also normal. And in the non-VOTC resection patients, both ipsilesional and contralesional ILF and IFOF also exhibited normal microstructural properties. Together, these results speak to the specificity of the damage as a result of surgery.

Abnormalities in the microstructural indices in circumscribed regions of the ipsilesional tracts are informative. AD has been shown to either increase (de Ruiter *et al*., 2011; Della Nave *et al*., 2011; Zhu *et al*., 2013; Counsell, 2006; Acosta-Cabronero *et al*., 2010) or decrease (Budde *et al*., 2009; Kubicki *et al*., 2013) as a result of WM pathology. Meanwhile, RD has been more consistently shown to increase due to demyelination (Song *et al*., 2002, 2005), and, specifically, myelin sheath degradation (Seal, *et al*., 2008, Klawiter *et al*., 2011). The results here showed increased AD values that might indicate axonal fragmentation, resulting in restricted diffusion along the principal tract direction, as well as increased RD values that might indicate myelin degradation, resulting in diffusion out of the WM bundle.

As noted previously, these circumscribed findings ought to be interpreted with caution in that there is a limited number of patients in the sample, and in some patients, the ipsilesional tracts could not be defined. Nevertheless, the results from the VOTC and the non-VOTC patients together reveal that these effects are likely a specific outcome of the resection without further degeneration of the WM as a result of surgery, and not due to a more pervasive WM deficits linked to epilepsy or other comorbidities.

#### 4.1.2 Normal microstructural indices in VOTC resection

Among the cases presented here, surprisingly, a single patient, OT, exhibited normal microstructural indices, even in the ipsilesional ILF and IFOF. This cannot be simply due to the age at surgery, as other patients were either younger (SN, UD) or older (KQ, NN) at the time of resection and none showed entirely normal microstructural indices. Similarly, the normal microstructure of the tracts cannot obviously be accounted for in terms of the time when the data were acquired postoperatively. One plausible explanation for the integrity of OT’s tracts is etiological — OT’s surgery was prompted by the presence of a dysembryoplastic neuroepithelial tumor (DNET). We speculate that the chronic growth of the DNET triggered reorganization in OT’s ipsilesional hemisphere such that the WM tracts were slowly displaced over time, and consequently, spared from damage due to the resection. However, we did not see this preservation in the only other patient with a DNET, UD, who evinced microstructural deficits ipsilesionally and network changes contralesionally. Of note, however, is that UD’s surgery occurred when he was 6 years old, but OT’s surgery took place when he was 13 years old. OT’s extended preoperative development may account for the difference between him and UD, but these speculative comments await additional experimental confirmation. We also consider UD in further detail below (Section 4.2.1).

### 4.2 Network properties of the contralesional hemisphere

In search for broader resection-induced changes, we characterized the connectivity of the contralesional occipito-temporal cortex by measuring transitivity and modularity (as these reflect segregation of the nodes into sub-networks) and characteristic path length and efficiency (as these reflect the ease of information flow in the network) (Bullmore and Sporns, 2009; Rubinov and Sporns, 2010; Paldino *et al*., 2019). There were no between-group differences across any of these dependent measures. However, on an individual level, while there were no changes to the network in the contralesional hemisphere of the LH VOTC resection patients or in the two non-VOTC resection patients, we did observe changes to the network in the contralesional hemisphere in the RH VOTC resection patients.

A similar pattern of network differences was reported in patients with LH lesions, previously (Crofts *et al*., 2011). While the etiology of the lesions in those patients – stroke – is different to the cases presented here, the hemispheric differences bear some resemblance. Also, Crofts et al. (2012) measured network communicability, a measure of the ease with which information can travel across a network. This is somewhat similar to the network efficiency that we measured here and for which we saw significant differences in the RH VOTC resection. The authors postulated that such alterations possibly arise from secondary degradation of long-range WM pathways that may or may not be directly connected to the damaged area.

#### 4.2.1 Differential hemispheric effects of resection on network properties

Given the small number of patients, we cannot definitively attribute the different profiles of network configuration to differences in hemispheric specialization. Notwithstanding the recommendation of caution in interpretation, the findings reported here are provocative enough to warrant some speculation, but additional exploration of this issue is obviously warranted.

There is well-known hemispheric lateralization of language to the LH and visual function to the RH in the majority of the population (Corballis, 1983). That we only see adverse changes to network properties in the two patients with RH VOTC resections might indicate that, in such cases, WM is reorganized in the contralesional LH to accommodate functions that are typically lateralized to, and dominant in, the RH. That is, the visual functions from the RH must be localized elsewhere postoperatively, and a potential location would be in the homologous LH regions. This transfer of function might be made possible via the corpus callosum, which plays a key role in plasticity after brain injury (Restani and Caleo, 2016) and whose maturation depends on coordinated activation across the two hemispheres (Pietrasanta *et al*., 2012).

One explanation of why there is no commensurate reorganization of the contralesional RH following LH VOTC resections is that the RH may already be finely tuned to subserve visual function. Remember that here, only the occipito-temporal regions were considered in the network topology. The cases of SN and TC support this view in that we do not see altered network properties in the contralesional RH when we know from fMRI studies that some visual functions of the resected regions have indeed emerged in the contralesional RH (as in the right VWFA in patients SN and TC, Table 1). The idea then is that, for visual function, the RH is already well tuned and can easily absorb the LH visual function, but the converse does not hold.

#### 4.2.2 Use of graph theory on structural data from pediatric epilepsy cases

Graph theory measures, applied to structural data, have previously been used in medial temporal lobe epilepsy in adults (Bonilha *et al*., 2012), where the authors showed normal network efficiency in patients prior to surgery. The absence of network changes pre-operatively suggests that epilepsy itself may not always alter the network efficiency in the epileptic brain, and that any changes are likely resection-induced. However, to our knowledge, aside from the present study, there has not been a systematic study of graph-theoretic measures applied to WM postoperative connectivity in children.

Because network properties rely on robust connectivity matrices, we studied the effects of using different threshold levels on the stability of the measures derived from tractography (see Supplementary Materials, Fig. D1). On the one hand, without thresholding (i.e. using data that potentially include spurious connections with low FA), only patients KQ and OT (right and left resection, respectively) had altered network properties. On the other hand, at an extremely high threshold, there were no alterations to the network properties, likely resulting in false negatives. Notably, in EK and DX (non-VOTC resection patients), the network properties of the contralesional temporo-occipital regions were consistently normal independent of threshold, suggesting that the alterations were likely a result of the resection to the VOTC and not due to the epilepsy per se.

Ultimately, using a reasonable threshold that was used to minimize noise, we found results from the RH VOTC resection cases that are consistent with results from a previous study that showed a moderate decrease in the network efficiency in epileptic patients (Taylor *et al*., 2018). It must be noted, however, that the result of Taylor et al. (2018) was based on an artificial resection of the (whole-brain) connectome data acquired preoperatively, whereas we provide an explicit characterization of the properties of the contralesional temporo-occipital network using in vivo data acquired postoperatively.

All that being said, we recognize that cognitive impairments in domains that are not directly associated with visual function (e.g. IQ below 70) might already be present premorbidly (Veersema *et al*., 2019) and might impact the network topography postoperatively. In our patients, only NN (with neurological comorbidities: extant polymicrogyria and low IQ) and DX showed some intermediate and high-level visual behavioral deficits (Liu, Freud *et al*., 2019). The other LH resection patients had no impairments and alterations of their contralesional network properties, while the RH VOTC resection patients had no impairments with alterations to their network properties. This heterogeneity confounds determination of the relationship between network properties vis-a-vis possible compensatory behavior. Nevertheless, network properties based on resting state data have already been shown to be reliable biomarkers for cognitive ability in children with epilepsy (Paldino *et al*., 2017b). Thus, it may be a worthwhile endeavor to determine whether network properties, as defined from structural connectivity, can be correlated with behavior in a larger sample of postoperative participants.

### 4.3 Normal visual perception despite persistent degradation in ipsilesional hemisphere

The main goal of this study was to determine the possible role of WM changes in relation to the preserved visual competence in pediatric patients with unilateral resections. A recent longitudinal study following individuals with anterior temporal lobectomy has documented changes over time to the FA of various WM tracts, including the bilateral ILF (Li *et al*., 2019). However, the authors did not measure behavioral outcomes in these patients. In Decramer et al (2019), recovery of behavioral function was seen in a single patient who had persistently damaged WM. As we only have cross-sectional diffusion data, albeit coupled with several behavior measures, we cannot comment on the longitudinal variations in the microstructural damage in our patients. However, the data from each patient used in the analyses were obtained at various times postoperatively, ranging from only a few months to more than a decade, and the patterns in microstructural damage are, nevertheless, largely similar across all the cases. It is, therefore, likely that the postoperative focal damage is persistent in the patients presented here and this is in the face of their largely normal higher-order perceptual abilities.

In sum, the current findings reveal an apparent dissociation between the postoperative structural damage and the intact behavioral competence in these patients. This finding contrasts with several studies reporting an association between compromised VOTC WM tracts and visual behavioral deficits, such as in prosopagnosia (Gomez *et al*., 2015, Thomas *et al*., 2009; Geskin and Behrmann, 2018). For instance, in the absence of obvious neurological damage and ample visual experience, individuals with prosopagnosia do not acquire normal face recognition skills. In such individuals, both the ILF and IFOF were abnormal and, interestingly, the severity of the face recognition deficit was correlated with the extent to which the ILF was compromised (Gomez *et al*., 2015, Thomas *et al*., 2009; Geskin and Behrmann, 2018). Along similar lines, there is also an association between perception of facial emotion and damage to the right IFOF (Philippi *et al*., 2009). Together, these findings demonstrate that WM atypicalities can have clear behavioral consequences. This is just not the case in many of the patients described here.

One explanation for the largely normal perceptual abilities, that we favor speculatively, is that the remaining intact cortex – the entire contralesional hemisphere together with the varying degree of intact tissue in the ipsilesional hemisphere in patients – is sufficient to support normal function and behavior. That is, the normal WM integrity outside the resection, without need for any enhancement either ipsi- or contralesionally, already allows for effective neural activity. The cases of SN and TC with LH VOTC resections offer strong support for this view. Whereas a typical visual word form area is localized in the LH in majority of the population, in SN and TC, this region emerged in the RH, and yet, we see no enhancement of the right WM pathways relative to controls. Additionally, the WM damage here is well-circumscribed even in the ipsilesional hemisphere. Whereas the degradation of WM associated with face processing deficits in neurodevelopmental cases is diffuse across a tract, the damage as a result of surgery in our patients is confined to the site of resection.

## 5 Conclusions

The goal of this study was to determine the neural basis of the largely normal perceptual abilities in children with unilateral resections of visual cortical regions as a result of pharmacoresistant epilepsy (save for patients NN with neurological comorbidities and DX). To do so, we adopted two complementary approaches to probe local and broader effects on the underlying WM. First, the microstructural indices, used to characterize specific fiber tracts, revealed focal damage only to the ipsilesional ILF and IFOF, without degradation of tracts distal to the resection. Moreover, the contralesional tracts exhibited normal microstructural indices indicating preserved WM integrity without enhancement. Second, a network approach uncovered changes to the contralesional network properties of only the patients with RH VOTC resection, but not in patients with LH VOTC resection nor in patients with non-VOTC resection. Because of the small number of participants, namely only two/four patients with RH/LH VOTC resection, respectively, this finding, although provocative, requires further confirmation. Taken together, we do not observe obvious patterns of reorganization of WM. Instead, our findings suggest that, in cases of unilateral resections to the VOTC in childhood, the category selectivity in the preserved contralesional hemisphere and the intact tissue in the ipsilesional hemisphere, together with structurally preserved ILF and IFOF outside the resection, might suffice to subserve normal visual perception.

## Acknowledgments

We thank the patients, the controls, and their families for their time and cooperation. We also thank Drs John Pyles and Tim Verstynen for assistance in determining the optimal scan protocols, as well as the MRI technologists Scott Kurdilla and Mark Vignone for help with the acquisition of the imaging data, and Dr Frank Yeh and the VisCog group at CMU for the fruitful discussions.

## Abbreviations

AD: axial diffusivity
DNET: dysembryoplastic neuroepithelial tumor
FA: fractional anisotropy
fMRI: functional magnetic resonance imaging
IFOF: inferior fronto-occipital fasciculus
ILF: inferior longitudinal fasciculus
LH: left hemisphere
MRI: magnetic resonance imaging
RD: radial diffusivity
RH: right hemisphere
ROI: region of interest
VOTC: ventral occipito-temporal cortex
WM: white matter

## References

1. Acosta-Cabronero, J., Williams, G. B., Pengas, G., and Nestor, P. J. Absolute diffusivities define the landscape of white matter degeneration in Alzheimer’s disease. Brain 2010; 133:529–539.

2. Arfanakis, K., Haughton, V. M., Carew, J. D., Rogers, B. P., Dempsey, R. J., and Meyerand, M. E. Diffusion tensor MR Imaging in diffuse axonal injury. American Journal of Neuroradiology 2002; 23:794–802.

3. Aung, W. Y., Mar, S., and Benzinger, T. L. Diffusion tensor MRI as a biomarker in axonal and myelin damage. Imaging in Medicine 2013; 5:427–440.

4. Behrmann, M. and Plaut, D. C. Distributed circuits, not circumscribed centers, mediate visual recognition. Trends in Cognitive Sciences 2013; 17:210–219.

5. Bonilha, L., Edwards, J. C., Kinsman, S. L., Morgan, P. S., Fridriksson, J., Rorden, C., Rumboldt, Z., Roberts, D. R., Eckert, M. A., and Halfor, J. J. Extrahippocampal gray matter loss and hippocampal deafferentation in patients with temporal lobe epilepsy. Epilepsia 2010; 51: 519–528.

6. Bonilha, L., Nesland, T., Martz, G. U., Joseph, J. E., Spampinato, M. V., Edwards, J. C., and Tabesh, A. Medial temporal lobe epilepsy is associated with neuronal fibre loss and paradoxical increase in structural connectivity of limbic structures. Journal of Neurology, Neurosurgery and Psychiatry 2012; 83:903–909.

7. Budde, M. D., Xie, M., Cross, A. H., and Song, S.K. Axial Diffusivity Is the Primary Correlate of Axonal Injury in the Experimental Autoimmune Encephalomyelitis Spinal Cord: A Quantitative Pixelwise Analysis. Journal of Neuroscience 2009; 29:2805–2813.

8. Bullmore, E. and Sporns, O. Complex brain networks: graph theoretical analysis of structural and functional systems. Nature Reviews Neuroscience 2009; 10:186–98.

9. Catani, M. Diffusion tensor magnetic resonance imaging tractography in cognitive disorders. Current Opinion in Neurology 2006; 19:599 –606.

10. Catani, M., Jones, D. K., Donato, R., ffytche, D. H. Occipito-temporal connections in the human brain. Brain 2003; 126:2093 –2107.

11. Corballis, M. C. (1983) Human Laterality. Auckland, New Zealand: Academic Press.

12. Counsell, S. J. Axial and Radial Diffusivity in Preterm Infants Who Have Diffuse White Matter Changes on Magnetic Resonance Imaging at Term-Equivalent Age. Pediatrics 2006; 117:376–386.

13. Crawford, J. R. and Howell, D. C. Comparing an individual’s test score against norms derived from small Samples. The Clinical Neurophsychologist 1998; 12:482–486.

14. Crofts, J. J., Higham, D. J., Bosnell, R., Jbabdi, S., Matthews, P. M., Behrens, T. E. J. Johansen-Berg, H. Network analysis detects changes in the contralesional hemisphere following stroke. NeuroImage 2011; 54:161–169.

15. Dale, A. M., Fischl, B., and Sereno, M. I. Cortical surface-based analysis: I. Segmentation and surface reconstruction. NeuroImage 1999; 9:179–194.

16. de Ruiter, M. B., Reneman, L., Boogerd, W., Veltman, D. J., Caan, M., Douaud, G., Lavini, C., Linn, S. C., Boven, E., van Dam, F. S. A. M., Schagen, S. B. Late effects of high-dose adjuvant chemotherapy on white and gray matter in breast cancer survivors: Converging results from multimodal magnetic resonance imaging. Human Brain Mapping 2011; 33:2971–2983.

17. Decramer, T., Premereur, E., Lagae, L., van Loon, J., Janssen, P., Sunaert, S., Theys, T. Patient MW: transient visual hemi-agnosia. Journal of Neurology 2019; 266:691–698.

18. Della Nave, R., Ginestroni, A., Diciotti, S., Salvatore, E., Soricelli, A., and Mascalchi, M. Axial diffusivity is increased in the degenerating superior cerebellar peduncles of Friedreich’s ataxia. Neuroradiology 2011; 53:367–372.

19. Dennis, E. L., Jin, Y., Kernan, C., Babikian, T., Mink, R., Babbitt, C., Johnson, J., Giza, C. C., Asarnow, R., Thompson, P. M. White matter integrity in traumatic brain injury: Effects of permissible fiber turning angle. In 2015 IEEE 12th International Symposium on Biomedical Imaging; pages 930–933. IEEE.

20. DeSalvo, M. N., Douw, L., Tanaka, N., Reisenberger, C., and Stufflebeam, S. M. Altered Structural Connectome in Temporal Lobe Epilepsy. Neuroradiology 2014; 270:842–848.

21. Destrieux, C., Fischl, B., Dale, A., and Halgren, E. Automatic parcellation of human cortical gyri and sulci using standard anatomical nomenclature. NeuroImage 2000; 53:1–15.

22. Fang-Cheng Yeh, Wedeen, V. J., and Tseng, W.-Y. I. Generalized q-sampling imaging. IEEE Transactions on Medical Imaging 2010; 29:1626–1635.

23. Geskin, J. and Behrmann, M. Congenital prosopagnosia without object agnosia? A literature review. Cognitive Neuropsychology 2018; 35:4–54.

24. Gomez, J., Pestilli, F., Witthoft, N., Golarai, G., Liberman, A., Poltoratski, S., Yoon, J., Grill-Spector, K. Functionally Defined White Matter Reveals Segregated Pathways in Human Ventral Temporal Cortex Associated with Category-Specific Processing. Neuron 2015; 85:216–227.

25. Helmstaedter, C., Beeres, K., Elger, C.E., Kuczaty, S., Schramm, J., Hoppe, C. (in press). Cognitive outcome of pediatric epilepsy surgery across ages and different types of surgeries: A monocentric 1-year follow-up study in 306 patients of school age. Seizure. doi: 10.1016/j.seizure.2019.07.021

26. Klawiter, E. C., Schmidt, R. E., Trinkause, K., Liang, H. F., Budde, M. D., Naismith, R. T., Song, S. K., Cross, A. H., Benzinger, T. L. Radial diffusivity predicts demyelination in ex vivo multiple sclerosis spinal cords. NeuroImage 2011; 55: 1454–1460.

27. Kubicki, M., Shenton, M. E., Maciejewski, P. K., Pelavin, P. E., Hawley, K. J., Ballinger, T., Swisher, T., Jabbar, G. A. Theremenos, H.W., Keshavan, M.S., Seidman, L.J., DeLisi, L.E. Decreased axial diffusivity within language connections: A possible biomarker of schizophrenia risk. Schizophrenia Research 2013; 148:67–73.

28. Li, W., An, D., Tong, X., Liu, W., Xiao, F., Ren, J., Niu, R., Tang, Y., Zhou, B., Lei, D., Jiang, Y., Luo, C., Yao, D., Gong, Q., Zhou, D. Different patterns of white matter changes after successful surgery of mesial temporal lobe epilepsy. NeuroImage: Clinical 2019; 21:10163.

29. Liu, T. T., Freud, E., Patterson, C., and Behrmann, M. Perceptual function and category-selective neural organization in children with resections of visual cortex. The Journal of Neuroscience 2019; 39: 6299–6314.

30. Liu, T. T., Nestor, A., Vida, M. D., Pyles, J. A., Patterson, C., Yang, Y., Yang, F. N., Freud, E., Behrmann, M. Successful reorganization of category-selective visual cortex following occipito-temporal lobectomy in childhood. Cell Reports 2018; 24:1113–1122.

31. Makris, N., Papadimitriou, G. M., Sorg, S., Kennedy, D. N., Caviness, V. S., and Pandya, D. N. The occipitofrontal fascicle in humans: A quantitative, in vivo, DT-MRI study. NeuroImage 2007; 37:1100–1111.

32. Mishkin, M., Ungerleider, L. G., and Macko, K. A. Object vision and spatial vision: two cortical pathways. Trends in Neurosciences 1983; 6:414–417.

33. Moen, K. G., Vik, A., Olsen, A., Skandsen, T., Haberg, A. K., Evensen, K. A. I., Eikenes, L. Traumatic Axonal Injury: Relationships Between Lesions in the Early Phase and Diffusion Tensor Imaging Parameters in the Chronic Phase of Traumatic Brain Injury. Journal of Neuroscience Research 2016; 635:623–635.

34. Ortibus, E., Verhoeven, J., Sunaert, S., Casteels, I., de Cock, P., and Lagae, L. Integrity of the inferior longitudinal fasciculus and impaired object recognition in children: A diffusion tensor imaging study. Developmental Medicine and Child Neurology 2011; 54:38–43.

35. Paldino, M. J., Chu, Z. D., Chapieski, M. L., Golriz, F., and Zhang, W. Repeatability of graph theoretical metrics derived from resting-state functional networks in pediatric epilepsy patients. British Journal of Radiology 2017a; 90:20160656.

36. Paldino, M. J., Golriz, F., Chapieski, M. L., Zhang, W., and Chu, Z. D. Brain network architecture and global intelligence in children with focal epilepsy. American Journal of Neuroradiology 2017b; 38:349–356.

37. Paldino, M. J., Golriz, F., Zhang, W., and Chu, Z. Normalization enhances brain network features that predict individual intelligence in children with epilepsy. PLoS ONE 2019; 14:e0212901.

38. Paldino, M. J., Hedges, K., Rodrigues, K. M., and Barboriak, D. P. Repeatability of quantitative metrics derived from MR diffusion tractography in pediatric patients with epilepsy. British Journal of Radiology 2014a; 87: 20140095.

39. Paldino, M. J., Zhang, W., Chu, Z. D., and Golriz, F. Metrics of brain network architecture capture the impact of disease in children with epilepsy. NeuroImage: Clinical 2017c; 13:201–208.

40. Philippi, C. L., Mehta, S., Grabowski, T., Adolphs, R., and Rudrauf, D. Damage to Association Fiber Tracts Impairs Recognition of the Facial Expression of Emotion. The Journal of Neuroscience 2009; 29:15089–15099.

41. Pietrasanta, M., Restani, L., and Caleo, M. The corpus callosum and the visual cortex: Plasticity is a game for two. Neural Plasticity 2012; 2012:838672.

42. Pustina, D., Doucet, G., Evans, J., Sharan, A., Sperling, M., Skidmore, C., Tracy, J. Distinct types of white matter changes are observed after anterior temporal lobectomy in epilepsy. PLoS ONE 2014; 9:e104211.

43. Restani, L. and Caleo, M. Reorganization of Visual Callosal Connections Following Alterations of Retinal Input and Brain Damage. Frontiers in Systems Neuroscience 2016; 10:86:1–17.

44. Rubinov, M. and Sporns, O. Complex network measures of brain connectivity: Uses and interpretations. NeuroImage 2010; 52:1059–1069.

45. Seal, M., Yucel, M., Fornito, A., Wood, S. J., Harrison, B. J., Walterfang, M., Pell, G. S., Pantelis, C. Abnormal white matter microstructure in schizophrenia: A voxelwise analysis of axial and radial diffusivity. Schizophrenia Research 2008; 1–3; 106–110.

46. Slinger, G., Sinke, M. R. T., Braun, K. P. J., Otte, W. M. White matter abnormalities at a regional and voxel level in focal and generalized epilepsy: A systematic review and meta-analysis. NeuroImage: Clinical 2016; 12: 902–909.

47. Song, S. K., Sun, S.W., Ramsbottom, M. J., Chang, C., Russell, J., and Cross, A. H. Dysmyelination revealed through MRI as increased radial (but unchanged axial) diffusion of water. NeuroImage 2002; 17:1429–1436.

48. Song, S.K., Yoshino, J., Le, T. Q., Lin, S.J., Sun, S.-W., Cross, A. H., Armstrong, R. C. Demyelination increases radial diffusivity in corpus callosum of mouse brain. NeuroImage 2005; 26:132–140.

49. Tavor, I., Yablonski, M., Mezer, A., Rom, S., Assaf, Y., and Yovel, G. Separate parts of occipito-temporal white matter fibers are associated with recognition of faces and places. NeuroImage 2014; 86:123–130.

50. Taylor, P. N., Han, C. E., Schoene-Bake, J.C., Weber, B., and Kaiser, M. Structural connectivity changes in temporal lobe epilepsy: Spatial features contribute more than topological measures. NeuroImage: Clinical 2015; 8:322–328.

51. Taylor, P. N., Sinha, N., Wang, Y., Vos, S. B., de Tisi, J., Miserocchi, A., McEvoy, A. W., Winston, G. P., Duncan, J. S. The impact of epilepsy surgery on the structural connectome and its relation to outcome. NeuroImage: Clinical 2018; 18:202–214.

52. Thomas, C., Avidan, G., Humphreys, K., Jung, K. J., Gao, F., and Behrmann, M. Reduced structural connectivity in ventral visual cortex in congenital prosopagnosia. Nature Neuroscience 2009; 12:29–31.

53. Veersema, T. J., van Schooneveld, M. M., Ferrier, C. H., van Eijsden, P., Gosselaar, P. H., van Rijen, P. C., Spliet, W. G. M., Muhlebner, A., Aronica, E. Braun, K. P. J. Cognitive functioning after epilepsy surgery in children with mild malformation of cortical development and focal cortical dysplasia. Epilepsy and Behavior 2019; 94:209–215.

54. Wakana, S., Caprihan, A., Panzenboeck, M. M., Fallon, J. H., Perry, M., Gollub, R. L., Hua, K., Zhang, J., Jiang, H., Dubey, P., Blitz, A., van Zijl, P., Mori, S. Reproducibility of quantitative tractography methods applied to cerebral white matter. NeuroImage, 2007; 36:630–644.

55. Wheeler-Kingshott, C. A. and Cercignani, M. About “axial” and “radial” diffusivities. Magnetic Resonance in Medicine 2009; 61:1255–1260.

56. Xu, J., Rasmussen, I.-A. J., Lagopoulos, J., and Haberg, A. Diffuse axonal injury in severe traumatic brain injury visualized using high-resolution diffusion tensor imaging. Journal of Neurotrauma 2007; 24:753–765.

57. Yeh, F.C., Verstynen, T. D., Wang, Y., Fernandez-Miranda, J. C., and Tseng, W.Y. I. Deterministic diffusion fiber tracking improved by quantitative anisotropy. PLoS ONE 2013; 8:e80713.

58. Zhang, W., Muravina, V., Azencott, R., Chu, Z. D., and Paldino, M. J. Mutual Information Better Quantifies Brain Network Architecture in Children with Epilepsy. Computational and Mathematical Methods in Medicine 2018; 2018:6142898.

59. Zhu, T., Zhong, J., Hu, R., Tivarus, M., Ekholm, S., Harezlak, J., Ombao, H., Navia, B., Cohen, R., Schifitto, G. Patterns of white matter injury in HIV infection after partial immune reconstitution: A DTI tract-based spatial statistics study. Journal of NeuroVirology 2013; 19:10–23.

