## Supplementary Materials for "Effects of unilateral cortical resection of the visual cortex on bilateral human white matter"

A. **Category-selective ROIs in pediatric cases of resection**

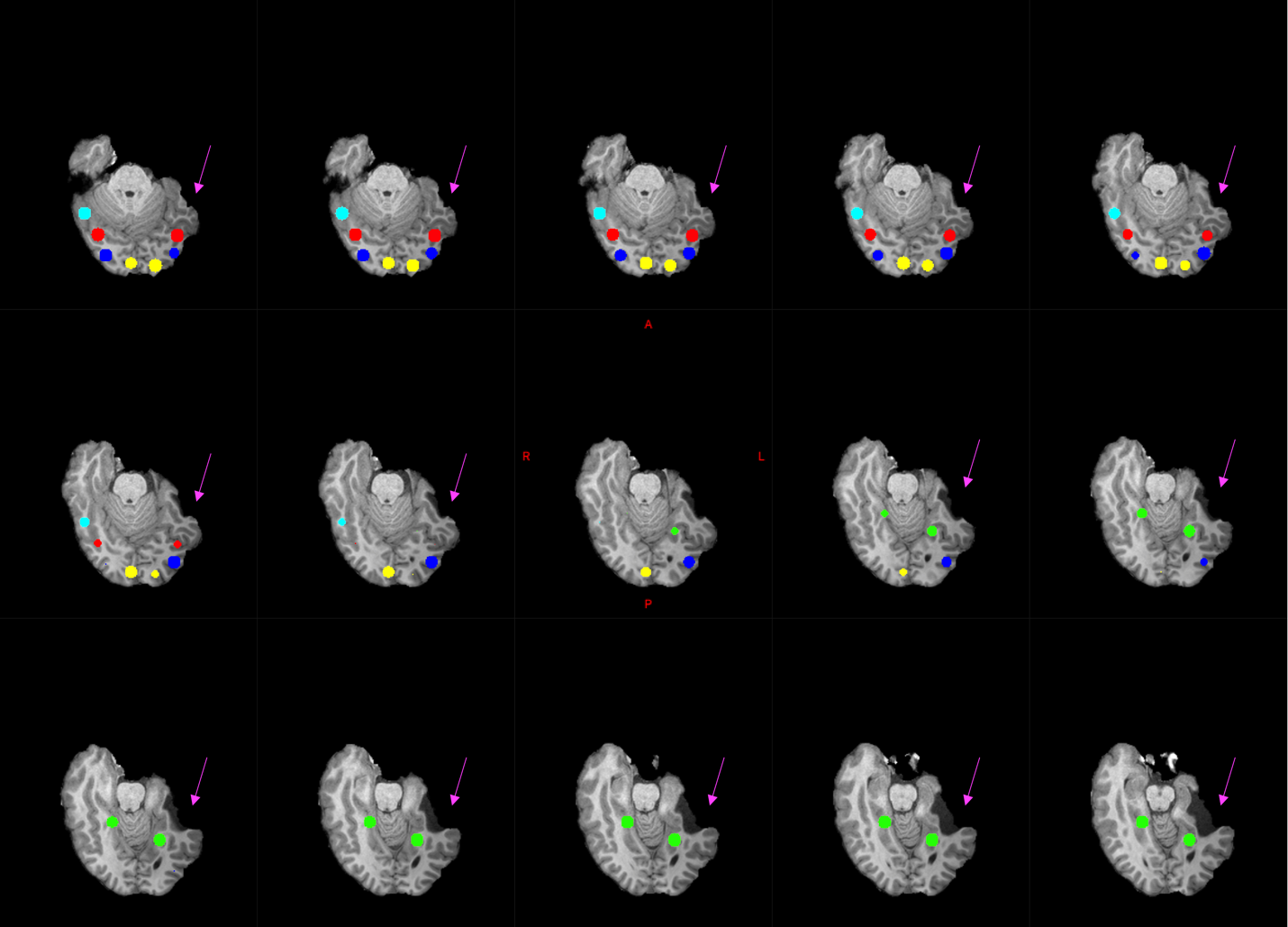

Figure A.1: Category-selective ROIs (Liu et al., 2018, 2019) were defined in Patient SN with left temporal lobe resection: Bilateral early visual cortex (yellow), fusiform face area (red), lateral occipital cortex (blue), parahippocampal place area (green), and unilateral visual word form area (VWFA, cyan). Magenta arrows indicate resected region. Note that the VWFA was localized to the right hemisphere.

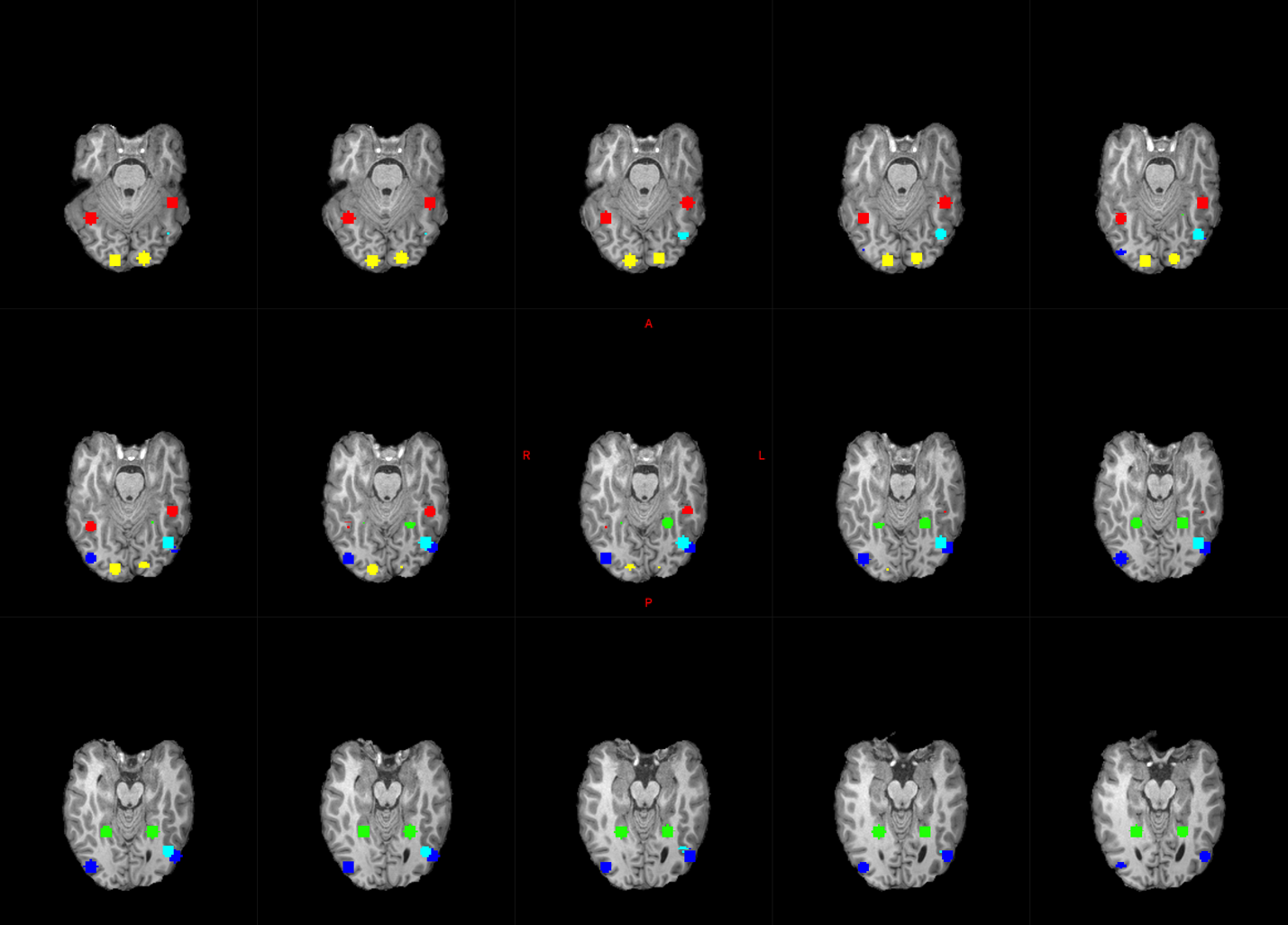

Figure A.2: Category-selective ROIs in Patient DX with left frontal resection (control without VOTC resection). Color scheme same as in Fig A.1.

B. **ROIs used as network nodes in network analysis**

In each hemisphere, 24 anatomical cortical ROIs, which were based on the Destrieux atlas (Destrieux *et al*., 2000) in FreeSurfer (Dale *et al*., 1999), were used as nodes in the network analysis of the contralesional hemisphere network in patients. These ROIs are listed below.

1. Gyrus and sulcus, inferior occipital

2. Gyrus, cuneus

3. Gyrus, middle occipital

4. Gyrus, superior occipital

5. Gyrus, lateral fusiform

6. Gyrus, lingual

7. Gyrus, parahippocampal

8. Gyrus, transverse superior temporal

9. Gyrus, lateral superior temporal

10. Gyrus, planum polare

11. Gyrus, planum temporale

12. Gyrus, inferior temporal

13. Gyrus, middle temporal

14. Occipital pole

15. Temporal pole

16. Sulcus, calcarine

17. Sulcus, middle and lunatus

18. Sulcus, superior and transversal occipital

19. Sulcus, anterior occipital

20. Sulcus, lateral occipito-temporal

21. Sulcus, medial and lingual

22. Sulcus inferior temporal

23. Sulcus, superior temporal

24. Sulcus, transverse temporal

C. **Behavioral experiments to assess visual perception**

Visual perception was assessed in all participants using a 14" laptop. Viewing distance was roughly 60 cm. The methods and results from most patients presented here (except SN and DX) were also presented in (Liu, Freud *et al*., 2019).

**C1 Global form perception**

Psychophysical thresholds were obtained from two separate tasks involving intermediate-level vision.

### **C1.1 Contour integration**

The contour integration task included two collinearity conditions (Hadad *et al*., 2010): target Gabor elements had either ±20° or ±0° collinearity, and participants pressed one of the two keys to indicate whether an embedded egg-like shape pointed rightwards or leftwards (Fig. C.1A). Background elements were varied according to a 1-up (after a wrong response), 3-down (after 3 correct responses) staircase procedure, and the experiment continued until 10 reversals in the staircase occurred. A practice session (10 trials) preceded each version of the task and the experiment took roughly 20 min. The threshold score was calculated from the geometrical mean spacing of the final 6 reversals. The active display (containing Gabor elements) extended about 17.6˚ horizontally and 12.6˚ vertically.

### **C1.2 Glass patterns**

Here, participants were instructed to press one of the two response keys to indicate which one of the two sequentially displayed glass patterns had more concentric “swirl” (Fig. C.1B). We varied the percentage of signal dots (Lewis *et al*., 2002) using a 1-up (after incorrect response), 3-down (after 3 correct responses) adaptive staircase method to measure the 75% threshold. The staircase started at 95% signal and terminated after 10 reversals, and the threshold was measured from the geometric mean of the last 6 reversals. The practice session contained 10 trials and the experiment took roughly 15 min. Each glass pattern was centrally presented and extended about 8.57˚ horizontally and 8.56˚ vertically.

**C2 Pattern recognition**

The integrity of high-level pattern recognition was explored using two tests, one for face and one for object recognition.

### **C2.1 Face recognition**

We used the Cambridge Face Memory Test for children (CFMT-C) (Croydon *et al*., 2014) (see Fig. C.1C) and followed the standard test instructions. Participants studied 5 faces and then, in subsequent trials, identified each ‘old’ face from amongst new, distractor faces. The test was conducted using upright and inverted faces in separate blocks. There were 60 trials in each orientation consisting of 15 introductory trials, 25 trials without noise, and 20 trials with added noise. Performance was the percent correct out of all 60 trials, separately for the upright and inverted version. Each face was presented centrally, extending about 3.4˚ horizontally and 4.9˚ vertically.

### **C2.2 Object recognition**

In this task (Fig. C.1D, adapted from Gauthier *et al.*, 1999), two objects were presented simultaneously — one above and one below the midline — for same/different discrimination. Each object subtended about 7.3˚ horizontally and 6.9˚ vertically on the screen. The task consisted of 100 trials, 40 same and 60 different (twenty per difference level), randomly intermixed. When the objects differed, they could differ at the basic (e.g., duck vs. vehicle), subordinate (e.g., chair vs. piano), or exemplar level (e.g., table1 vs. table2), reflecting increasing perceptual similarity. The display remained on the screen until response, with one key indicating ‘same’ and another ‘different’. Instructions encouraged both speed and accuracy (and both were measured), and a 25-trial practice block familiarized the participant with the speeded task.

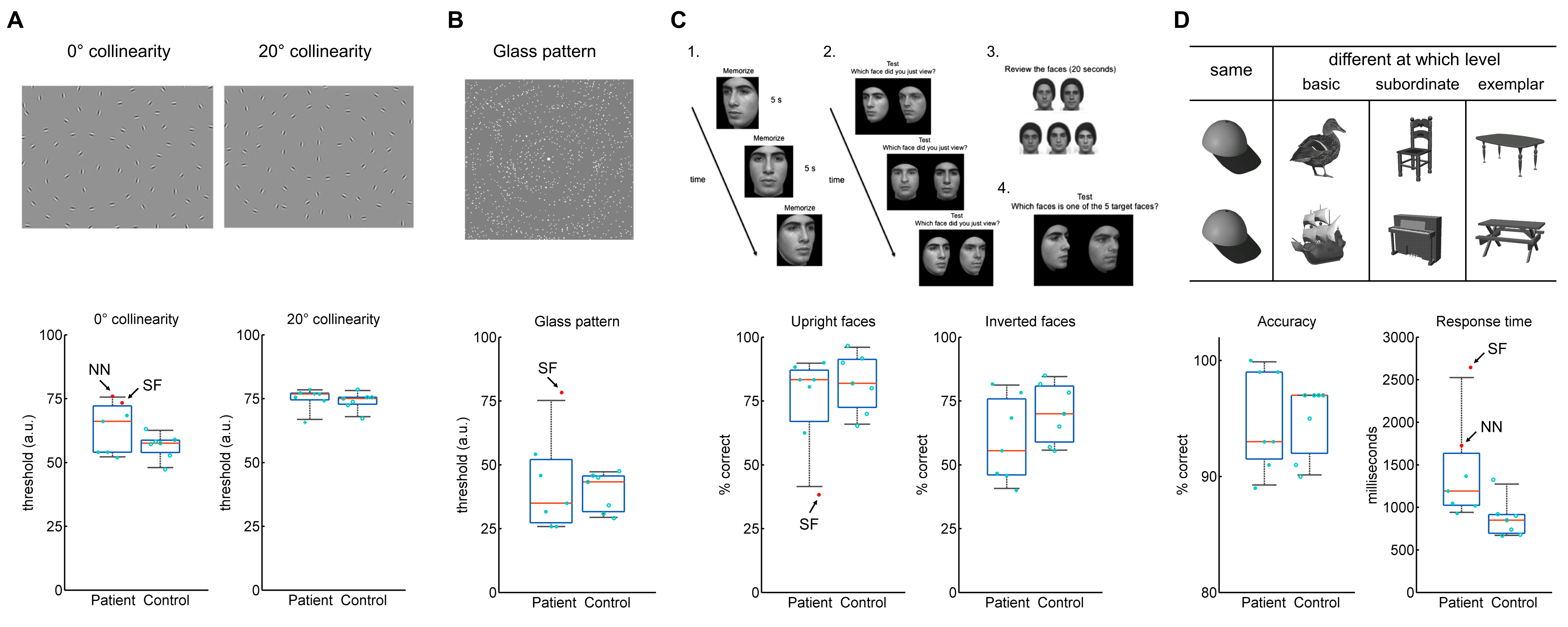

Figure C.1: Experimental design and results of the behavioral experiments. Examples of stimuli for each experiment are shown. **A**. Global form. Task: Participants viewed a brief presentation and indicated the leftward or rightward pointing of the embedded “egg-like” shape. Results: patients and controls were found to have similar thresholds, except that the threshold in the ±0° collinearity condition in patient NN was outside the control range. **B**. Glass pattern (Lewis *et al*., 2002). Task: Participants pressed a button to indicate which of the two displays had a more concentric swirl. Results: All patients showed normal threshold. **C**. CFMT-C (Croydon *et al*., 2014). Task: Participants were instructed to remember target faces and subsequently identify them amongst an array of distractor faces. Results: All patients showed normal face recognition abilities. **D**. Object recognition experiment (Gauthier *et al*., 1999). Task: Participants made same/different discriminations on pairs of objects and pressed a “same” or “different” button to indicate their response. Results: Patients and controls were found to have similar accuracy and RTs, excluding patient NN who exhibited slower RT.

D. **Effects of using different thresholds in binarizing the connectivity matrices**

We generated a connectivity matrix in DSI Studio, such that, for any ROI pair, the connectivity value was the mean FA of all tracts that pass through both ROIs. We binarized the connectivity matrix for each individual based on the mean (m) and standard deviation (s) values of all FA>0.
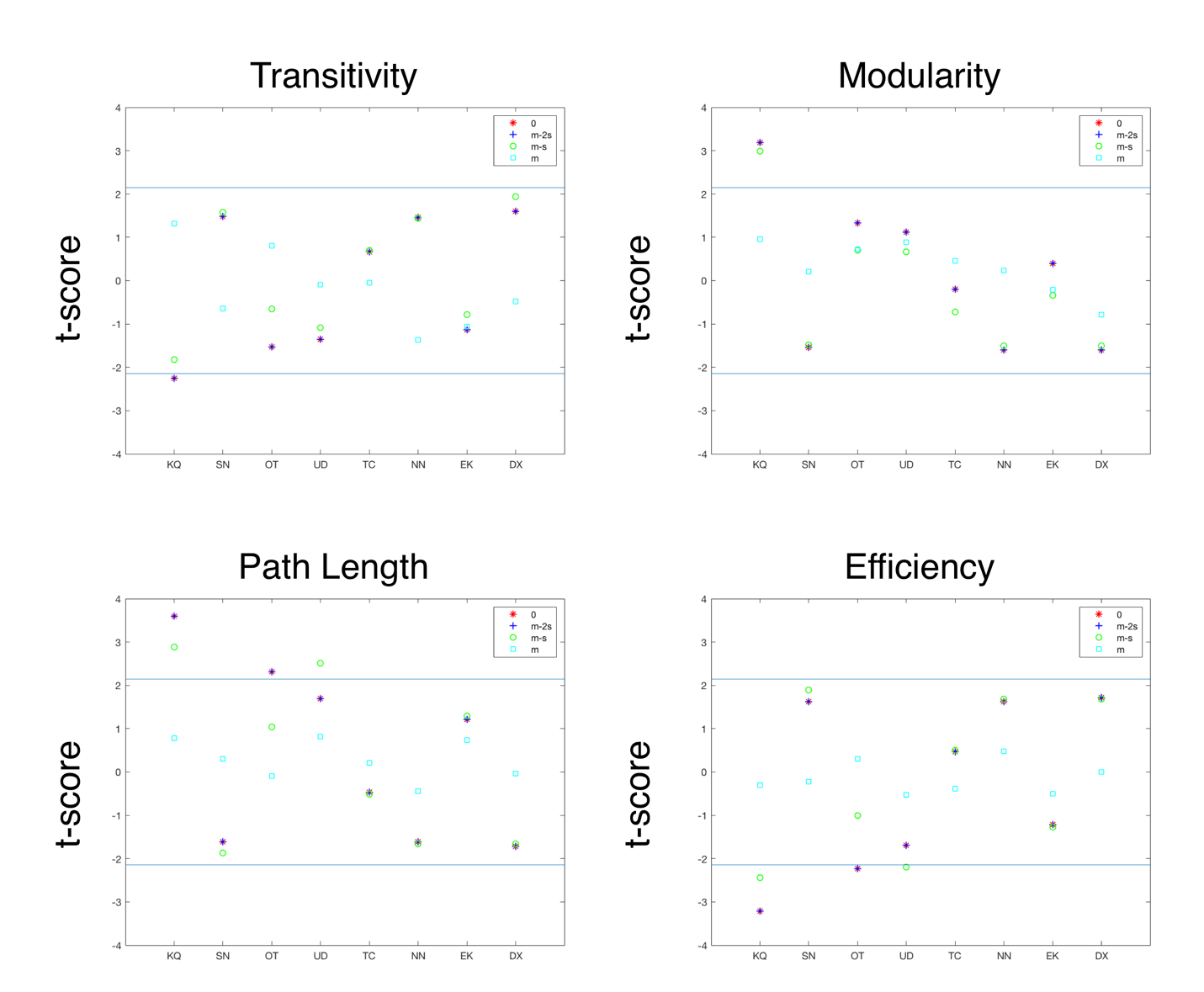

Figure D.1: Effects of using different thresholds on the network properties. At a higher threshold (m) there were no significant differences between any of the patients and controls in any of the dependent measures – i.e. all t-scores were within normal range (blue lines). At lower thresholds (0 and m-2s), only one patient with left resection, OT, had a significantly different path length and efficiency (all |t|>2.14, all p<0.05), and in the two patients with right resection, KQ had significantly different properties to controls, while these effects in UD disappeared. m: mean of

**References**

Croydon, A., Pimperton, H., Ewing, L., Duchaine, B. C., Pellicano, E. The Cambridge Face Memory Test for Children (CFMT-C): a new tool for measuring face recognition skills in childhood. Neuropsychologia 2014; 62:60–67.

Dale, A. M., Fischl, B., and Sereno, M. I. Cortical surface-based analysis: I. Segmentation and surface reconstruction. NeuroImage 1999; 9:179–194.

Destrieux, C., Fischl, B., Dale, A., and Halgren, E. Automatic parcellation of human cortical gyri and sulci using standard anatomical nomenclature. NeuroImage 2000; 53:1–15.

Gauthier, I., Behrmann, M., Tarr, M. J. Can face recognition really be dissociated from object recognition? Journal of Cognitive Neuroscience 1999; 11:349–370.

Hadad, B., Maurer, D., and Lewis, T. L. The effects of spatial proximity and collinearity on contour integration in adults and children. Vision Research 2010; 50:772–778.

Lewis T. L., Ellemberg, D., Maurer, D., Wilkinson, F., Wilson, H. R., Dirks, M., et al. Sensitivity to global form in glass patterns after early visual deprivation in humans. Vision Research 2002; 42:939–948.

Liu, T. T., Freud, E., Patterson, C., and Behrmann, M. June 5, 2019. Perceptual function and category-selective neural organization in children with resections of visual cortex. The Journal of Neuroscience 10.1523/JNEUROSCI.3160-18.2019
